# RaptorX-Single: single-sequence protein structure prediction by integrating protein language models

**DOI:** 10.1101/2023.04.24.538081

**Authors:** Xiaoyang Jing, Fandi Wu, Xiao Luo, Jinbo Xu

**Affiliations:** MoleculeMind Ltd., Beijing, China; Institute of Computing Technology, Chinese Academy of Sciences, Beijing, China; Toyota Technological Institute at Chicago, Chicago, IL, USA; Shanghai Artificial Intelligence Laboratory, Shanghai, China

**Keywords:** Protein structure prediction, protein language model, single sequence protein structure prediction, antibody structure prediction, single mutation effect

## Abstract

Protein structure prediction has been greatly improved by deep learning in the past few years. However, the most successful methods rely on multiple sequence alignment (MSA) of the sequence homologs of the protein under prediction. In nature a protein folds in the absence of its sequence homologs and thus, a MSA-free structure prediction method is desired. Here we develop a single sequence-based protein structure prediction method RaptorX-Single by integrating several protein language models and a structure generation module and then study its advantage over MSA-based prediction methods. Our experimental results indicate that in addition to running much faster than MSA-based methods such as AlphaFold2, RaptorX-Single outperforms AlphaFold2 and other MSA-free methods in predicting the structure of antibodies, proteins of very few sequence homologs and single mutation effects. RaptorX-Single also compares favorably to MSA-based AlphaFold2 when the protein under prediction has a large number of sequence homologs.

## Introduction

In the past few years computational protein structure prediction has been revolutionized by the application of coevolution information and deep learning, as evidenced first by RaptorX [1,2] and then AlphaFold2 [3] and other similar methods. These successful methods make use of sequence homologs of the protein under prediction and coevolution information derived from their multiple sequence alignment (MSA). Although very powerful, these methods are not perfect. For example, in nature a protein folds without knowledge of its sequence homologs, so predicting protein structure based upon some non-natural conditions such as MSAs (and coevolution information) does not reflect very well how a protein actually folds. It also takes much time to search for the sequence homologs of a protein, especially considering that the sequence databases are growing rapidly. Further, the MSA-based methods do not perform well on small-sized protein families. Experimental results show that MSA-based methods do not fare well on flexible regions such as loop regions [4] and highly variable antibody CDR regions. Further, MSA are not sensitive for sequence mutations, the MSA based methods may not fare well on the prediction of single-point mutational effects [5].

To reduce the dependence of protein structure prediction on MSA, we have studied protein structure prediction without using coevolution information in Xu et al [6], and demonstrated that in the absence of coevolution information, deep learning can still predict more than 50% of the most challenging CASP13 targets. This implies that deep learning indeed has captured sequence-structure relationships useful for tertiary structure prediction. Nevertheless, in this work we still used sequence profiles derived from sequence homologs. In the past few months, several groups [7,8,7,9–11] have studied deep learning methods for single-sequence based protein structure prediction by making use of protein language models. Although running much faster, on average these methods are less accurate than MSA-based AlphaFold2.

To further study the advantage of single-sequence based methods, we have developed RaptorX-Single for single sequence-based protein structure prediction. Our methods take an individual protein sequence as input and then feed it into protein language models to produce sequence embedding, which is then fed into a modified Evoformer module and a structure generation module to predict atom coordinates. Differing from other single sequence methods, our methods use a combination of three well-developed protein language models [12–14] instead of only one. Experimental results show that our RaptorX-Single not only runs much faster than MSA-based AlphaFold2, but also outperforms it on antibody structure prediction, orphan protein structure prediction and single mutation effect prediction. RaptorX-Single also exceeds other single-sequence based methods and compares favorably to AlphaFold2 when the protein under prediction has a large number of sequence homologs.

## Method

### Protein language models

Protein language models have been developed to model individual protein sequences. These models mainly use attention-based deep neural networks to capture long-range inter-residue relationships. Here we make use of three pretrained protein language models including ESM-1b [12], ESM-1v [13] and ProtTrans [14]. Meanwhile, ESM-1b is a Transformer of ∼650M parameters that was trained on UniRef50 [15] of 27.1 million protein sequences. ESM-1v employs the same network architecture as ESM-1b, but was trained on Uniref90 with 98 million protein sequences. For ProtTrans, we use the ProtT5-XL model of 3 billion parameters that was trained on a newer UniRef50 of 45 million sequences.

### Network architecture

The overall network architecture of our method is shown in Figure 1, which mainly consists of three modules: the sequence embedding module, the modified Evoformer module and the structure generation module. Given an individual protein sequence as input, the embedding module generates the sequence embedding of the input sequence and its pair representations, by making use of one or three protein language models. In the embedding module, the one-hot encoding of the input sequence passes through a linear layer to generate the initial sequence embedding, then it combines the sequence representation from protein language models to create a new sequential embedding. The initial pair embedding is generated by adding two primary sequence embedding (rowwise and columnwise), and it then combines the attention maps from the last two layers of protein language models to create a new pairwise embedding. We also add the relative positional encoding in the pairwise embedding.

**Figure 1.**
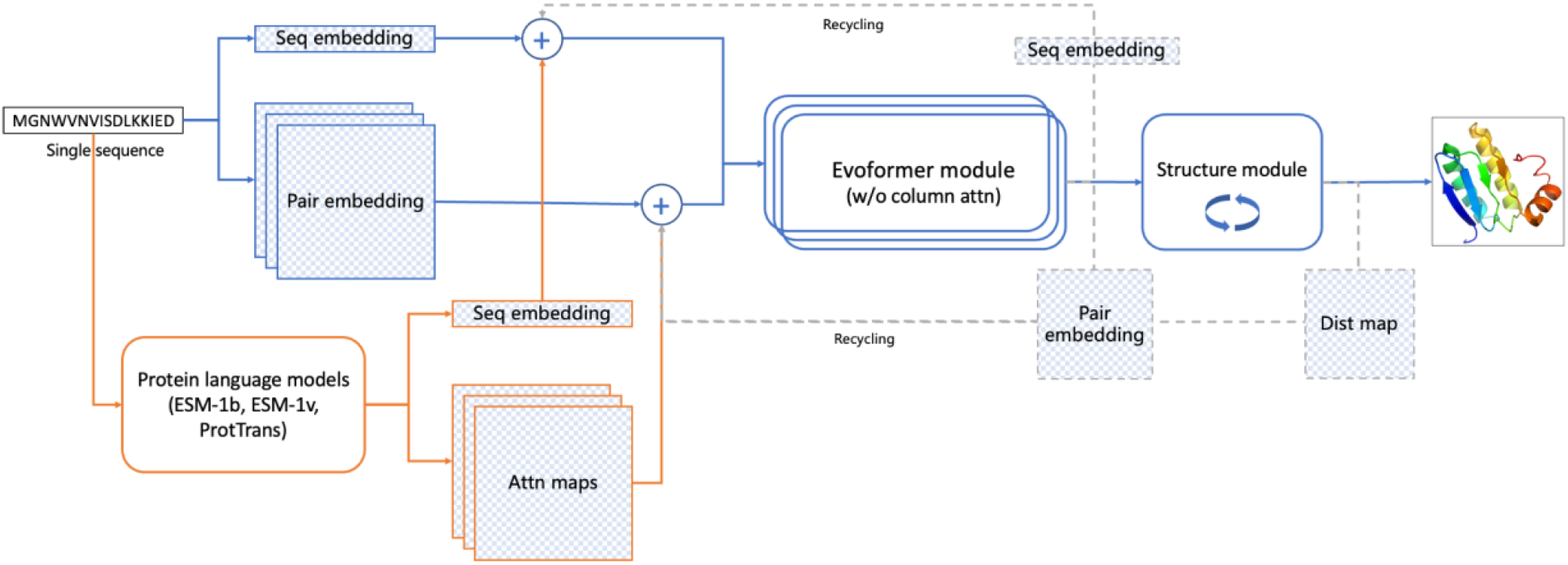
Our deep network architecture for single sequence-based protein structure prediction.

The sequence and pair embedding are updated iteratively in the Evoformer module consisting of 24 modified Evoformer layers. Our Evoformer differs from the original one in AlphaFold2 in that ours does not have the column self-attention layer, which is meaningless for an individual sequence.

The structure module is similar to that of AlphaFold2, which mainly consists of 8 IPA (invariant point attention) and transition layers with shared weights. Our structure module is different in that we use a linear layer to integrate the scalar, point, and pair attention values in the IPA model while AlphaFold2 uses only addition. The structure module outputs both the predicted atom 3D coordinates and confidence scores (i.e. pLDDT).

### Training and test data

#### Training data

The training data consists of ∼340k proteins. Among them, there are 80852 proteins of different sequences that have experimental structures released before January 2020 in PDB (denoted as BC100). We cluster the proteins in BC100 by 40% sequence identity and denote the clustering result as BC100By40. The remaining 264k proteins have tertiary structures predicted by AlphaFold2 (denoted as distillation data). The proteins in the distillation dataset are extracted from Uniclust30_2018_08 [16] and share no more than 30% sequence identity. During each training epoch, one protein is randomly sampled from each BC100By40 cluster to form a set of training proteins, with the acceptance rate determined by the sequence length (0.5 for lengths under 256, 0.5-1 for lengths between 256 and 512, and 1.0 for lengths over 512). In each epoch proteins are also sampled from the distillation data by the ratio of 1:3 between BC100By40 and the distillation data.

#### Antibody data for fine-tuning

In order to improve the performance on antibody targets, we construct an antibody training set for fine-tuning. Specifically, we use experimental structures from SAbDa [17] released before 2021/03/31 as training samples and split each target into chains. In total, the training set contains 5033 heavy and light chains. The validation set consists of 178 antibody structures released between 2021/04/01 and 2021/06/30.

#### Training procedure

The model was implemented using pytorch [18], and the distributed training on multi-GPUs was based on pytorch-lightning [19]. We used the AdamW [20] optimizer with β1=0.9, β2=0.999, ε=10−8 and weight decay of 0.0001 for all models. Over the first ∼1000 steps, the learning rate warmed up linearly from 1e-6 to 5e-4, remained at 5e-4 for the first 1/3 training steps, and then linearly decreased to 1e-6 for the remaining 2/3 training steps. The model is initially trained on protein crops of size 256 for the first 2/3 of the training steps, and then trained on crop sizes of 384 for the remaining 1/3 of the training steps. The training losses include pairwise losses and structure losses. The pairwise losses are distogram distance and orientation loss, as employed in trRosetta [21]. The structure losses include the Frame Aligned Point Error (FAPE) [3] loss with a clamp of 20 Angstrom and the pLDDT loss. To improve the performance of the model, we also implemented the recycling strategy during training. The number of recycling iterations is randomly sampled from 0 to 3. Each model is trained on 32 GPUs with accumulated gradients 4, thus the batch size is 128.

In total, we have trained four models with 150 epochs by combining the protein language models in different ways. RaptorX-Single (1b), RaptorX-Single (1v) and RaptorX-Single (pt) make use of ESM-1b, ESM-1v and ProtTrans respectively, while RaptorX-Single makes use of the three protein language models together. In the fine-tuning stage for antibody, we fine-tuned all four models 50 epochs with the learning rate linearly decreasing from 2e-4 to 1e-5, and obtained the corresponding four models for antibody structure prediction: RaptorX-Single-Ab (1b), RaptorX-Single-Ab (1v), RaptorX-Single-Ab (pt), and RaptorX-Single-Ab.

### Evaluation

#### Benchmark datasets

We evaluated our methods on three antibody datasets (SAbDab-Ab, IgFold-Ab and nanobody), one orphan protein dataset and one single mutation effects dataset.

- **IgFold-Ab dataset:** It includes 67 non-redundant antibodies from the IgFold work [22] that were released after July 1, 2021.
- **SAbDab-Ab dataset:** It is a non-redundant dataset (sequence identity lower than 95%) derived from SAbDab [17], which has 202 antibodies released in the first 6 months of 2022.
- **Nanobody dataset:** It includes 60 nanobodies of different sequences derived from SAbDab released in the first 6 months of 2022.
- **Orphan dataset:** It consists of 11 proteins released between Jan 01, 2020 and April 12, 2022. The proteins in this set do not have any sequence homologs in BFD [3], MGnify [23], Uniref90 [15] and Uniclust30 [16]. That is, they are not similar to any proteins used for training the protein language models and our deep learning models.
- **Rocklin dataset**. The Rocklin dataset has 14 native and de novo designed proteins and their stability measures of 10,674 single mutations. The stability was evaluated using thermal and chemical denaturation. The source data is from Rocklin et al [24] and here we use the data collected by Strokach et al [25].

#### Performance metrics

For antibodies, we evaluate the backbone root-mean-squared-deviation (RMSD) of the framework (Fr) and CDR (CDR-1, CDR-2, CDR-3) regions of the heavy and light chains separately. The CDR regions are classified using ANARCI, and the backbone RMSD is calculated by PyRosetta [26]. For orphan targets, we evaluate the quality of the predicted structures by TMscore [27], global distance test–total score (GDT_TS) and global distance test–high accuracy (GHT_HA) [28]. For the single mutation effects prediction, we calculate the Pearson correlation coefficient (PCC) between the predicted structure changes and the stability data. The structure change is measured by the TMscore of predicted structures between the wild and mutant sequences.

#### Baseline methods

We compared the performance of our methods with MSA-based AlphaFold2 and three single sequence-based methods (ESMFold [7], OmegaFold [8] and HelixFold-Single [11]). For AlphaFold2, we search the Uniclust30 [16] as of August 2018, Uniref90 [15] as of January 2020 and Metaclust [23] as of December 2018 and BFD [3] using HHblits and Jackhmmer to build its MSA. Additionally, we also report the performance of AlphaFold2 using a single sequence as input. For the antibody datasets, we also compare our method with antibody-specific methods DeepAb [29] and IgFold [22].

## Results

### Performance on the IgFold-Ab dataset

As shown in Table 1, our antibody-specific method RaptorX-Single-Ab consistently outperforms other methods for antibody structure prediction, especially in the CDR H3 region as shown in Figure 2 (A). The MAS-based AlphaFold2 performs comparably with other single sequence methods (HelixFold-Single, OmegaFold and ESMFold), but is not as good as the antibody-specific methods DeepAb and IgFold. AlphaFold2 (Single) performs much worse than AlphaFold2 (MSA), mainly because AlphaFold2 was trained on MSAs instead of single sequences. Our antibody-specific methods outperform our non-fine-tuned methods, suggesting the value of the fine-tuning phase on antibody data.

**Table 1.** The average RMSD of predicted tertiary structures for the IgFold-Ab dataset.

|  | RMSD (H) |  |  |  | RMSD (L) |  |  |  |
| --- | --- | --- | --- | --- | --- | --- | --- | --- |
|  | Fr | CDR-1 | CDR-2 | CDR-3 | Fr | CDR-1 | CDR-2 | CDR-3 |
| AlphaFold2 (MSA) | 0.48 | 0.77 | 0.76 | 3.55 | 0.43 | 0.96 | 0.45 | 1.26 |
| AlphaFold2 (Single) | 10.84 | 15.34 | 15.48 | 16.33 | 8.98 | 13.54 | 16.13 | 15.14 |
| HelixFold-Single | 0.56 | 0.85 | 0.95 | 5.01 | 0.51 | 1.10 | 0.57 | 1.60 |
| OmegaFold | 0.47 | 0.75 | 0.74 | 3.70 | 0.41 | 0.93 | 0.43 | 1.35 |
| ESMFold | 0.51 | 0.84 | 0.91 | 4.10 | 0.43 | 1.16 | 0.52 | 1.44 |
| DeepAb | 0.43 | 0.80 | 0.74 | 3.28 | 0.38 | 0.86 | 0.45 | 1.11 |
| IgFold | 0.45 | 0.80 | 0.75 | 2.99 | 0.45 | 0.83 | 0.51 | 1.07 |
| RaptorX-Single | 0.51 | 0.86 | 0.90 | 4.33 | 0.46 | 1.13 | 0.54 | 1.95 |
| RaptorX-Single-Ab | 0.38 | 0.63 | 0.60 | 2.65 | 0.35 | 0.69 | 0.39 | 0.88 |

**Figure 2.**
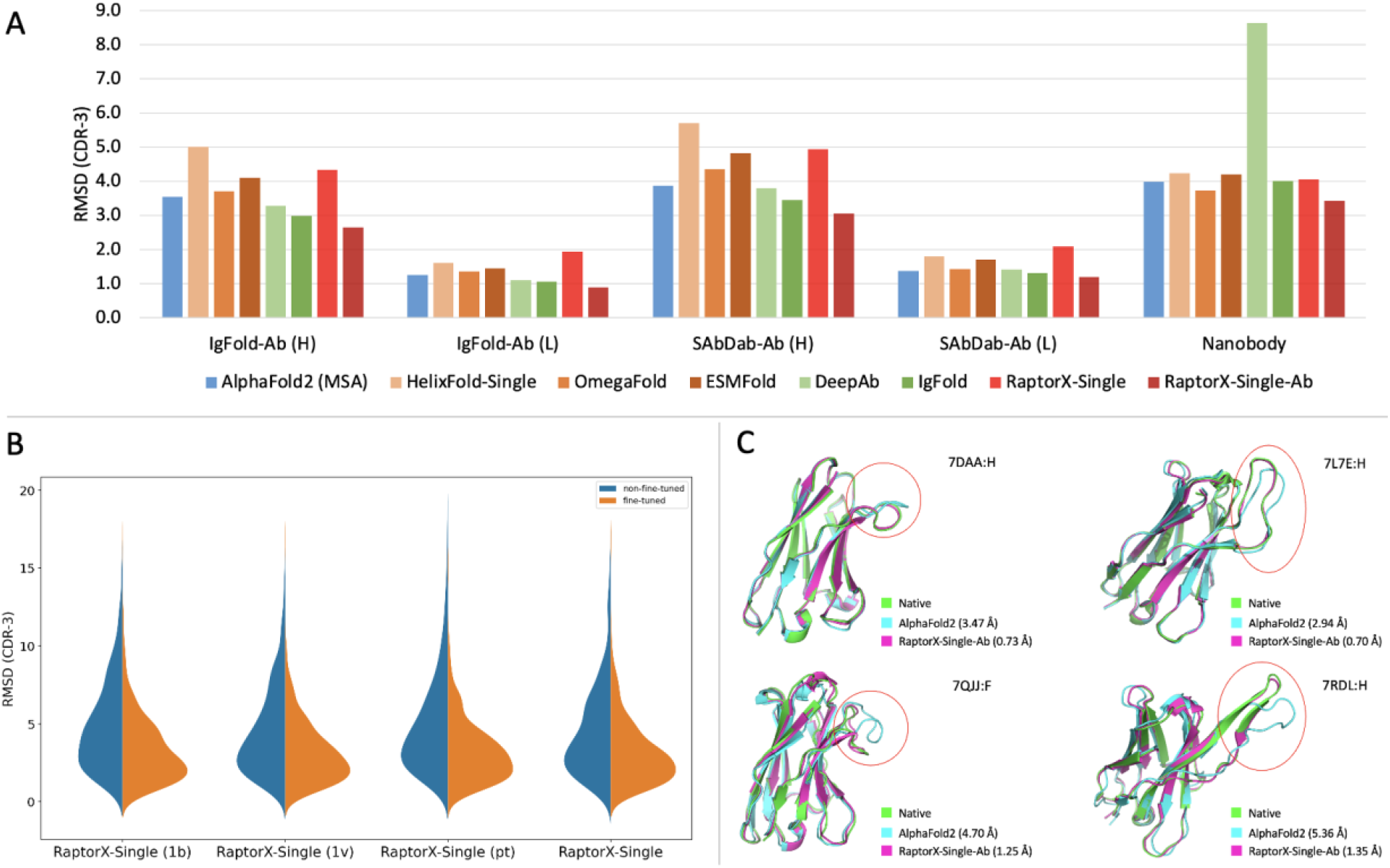
Performance comparison of various methods on antibody structure prediction. (A) The average RMSD of predicted CDR-3 structures on different datasets. (B) The CDR-3 RMSD distribution of our non-fine-tuned and antibody fine-tuned models by combining protein language models in different ways. The antibody fine-tuned methods consistently outperform the non-fine-tuned methods (C) Comparison of RaptorX-Single-Ab and MSA-based AlphaFold2 on the CDR H3 structure prediction (marked by red circle) for targets 7DAA, 7L7E, 7QJJ and 7RDL.

### Performance on the SAbDab-Ab dataset

Our fine-tuned antibody-specific method greatly outperforms other methods regardless of metrics, as shown in Table 2. AlphaFold2 (MSA) performs slightly better than other single sequence methods (HelixFold-Single, OmegaFold and ESMFold), and is comparable with the antibody-specific methods DeepAb and IgFold. Our fine-tuned antibody-specific methods outperform our non-fine-tuned methods, while there is no obvious difference between models using different protein language models as shown in Table S2. As shown in Figure 2 (C), our RaptorX-Single-Ab significantly outperforms AlphaFold2 (MSA) on some targets.

**Table 2.** The average RMSD of predicted tertiary structures for the SAbDab-Ab dataset.

|  | RMSD (H) |  |  |  | RMSD (L) |  |  |  |
| --- | --- | --- | --- | --- | --- | --- | --- | --- |
|  | Fr | CDR-1 | CDR-2 | CDR-3 | Fr | CDR-1 | CDR-2 | CDR-3 |
| AlphaFold2 (MSA) | 0.63 | 1.08 | 0.89 | 3.82 | 0.59 | 0.89 | 0.69 | 1.39 |
| AlphaFold2 (Single) | 8.85 | 12.30 | 11.59 | 15.24 | 8.82 | 13.28 | 15.13 | 14.62 |
| HelixFold-Single | 0.71 | 1.15 | 1.10 | 5.50 | 0.66 | 1.10 | 0.79 | 1.84 |
| OmegaFold | 0.63 | 1.05 | 0.86 | 4.11 | 0.58 | 0.90 | 0.69 | 1.42 |
| ESMFold | 0.64 | 1.11 | 1.02 | 4.56 | 0.60 | 1.16 | 0.72 | 1.74 |
| DeepAb | 0.62 | 1.08 | 0.90 | 3.83 | 0.66 | 0.96 | 0.75 | 1.43 |
| IgFold | 0.66 | 1.15 | 0.95 | 3.65 | 0.65 | 0.96 | 0.80 | 1.40 |
| RaptorX-Single | 0.64 | 1.17 | 1.06 | 4.66 | 0.64 | 1.12 | 0.77 | 2.14 |
| RaptorX-Single-Ab | 0.57 | 1.01 | 0.82 | 3.24 | 0.53 | 0.79 | 0.66 | 1.24 |

### Performance on the Nanobody dataset

Nanobody is an increasingly popular modality for therapeutic development [30]. Compared with traditional antibodies, nanobodies lack a second immunoglobulin chain and have increased CDR3 loop length, which makes the nanobody structure prediction challenging. As shown in Table 3, no methods perform very well on the CDR-3 region, but our fine-tuned antibody-specific models still outperform other methods. AlphaFold2 (MSA) outperforms HelixFold-Single and ESMFold but underperforms OmegaFold. DeepAb performs poorly on nanobodies, possibly because it is trained for paired antibody structure prediction. RaptorX-Single-Ab performs the best, demonstrating the value of combining three protein language models.

**Table 3.** The average RMSD of the predicted structures for the Nanobody dataset.

|  | RMSD |  |  |  |
| --- | --- | --- | --- | --- |
|  | Fr | CDR-1 | CDR-2 | CDR-3 |
| AlphaFold2 (MSA) | 0.73 | 2.05 | 1.15 | 4.01 |
| AlphaFold2 (Single) | 9.3373 | 12.6722 | 12.3947 | 17.8738 |
| HelixFold-Single | 0.86 | 1.99 | 1.18 | 4.20 |
| OmegaFold | 0.71 | 2.02 | 1.12 | 3.77 |
| ESMFold | 0.80 | 2.06 | 1.12 | 4.23 |
| DeepAb | 0.92 | 2.38 | 1.34 | 8.76 |
| IgFold | 0.82 | 1.93 | 1.29 | 4.27 |
| RaptorX-Single | 0.83 | 2.19 | 1.14 | 4.06 |
| RaptorX-Single-Ab | 0.82 | 1.78 | 1.06 | 3.50 |

### Performance of different combinations of protein language models

We have trained four models by combining the protein language models in different ways. RaptorX-Single (1b), RaptorX-Single (1v) and RaptorX-Single (pt) make use of ESM-1b, ESM-1v and ProtTrans respectively, while RaptorX-Single makes use of the three protein language models together. The detailed performance of different protein language model combinations are shown in Table S1-4, and Figure 2 (B) shows the performance on CDR-3 regions. Overall, by combining three protein language models, RaptorX-Single and RaptorX-Single-Ab performs better than those that use only a single protein language model.

### Performance on the Orphan dataset

Orphan proteins lack evolutionary homologs in structure and sequence databases and thus, are very challenging for MSA-based methods. Here we construct an orphan protein dataset, in which no protein has any sequence homologs in the 4 widely-used sequence databases BFD, MGnify, Uniref90 and Uniclust30. As shown in Table 4, almost all single sequence methods outperform AlphaFold2 on this dataset and our method RaptorX-Single achieved the best performance. However, neither MSA nor single sequence methods can predict the correct fold of most orphan proteins, mainly because current single sequence methods are still implicitly making use of homologous information learned by protein language models.

**Table 4.**
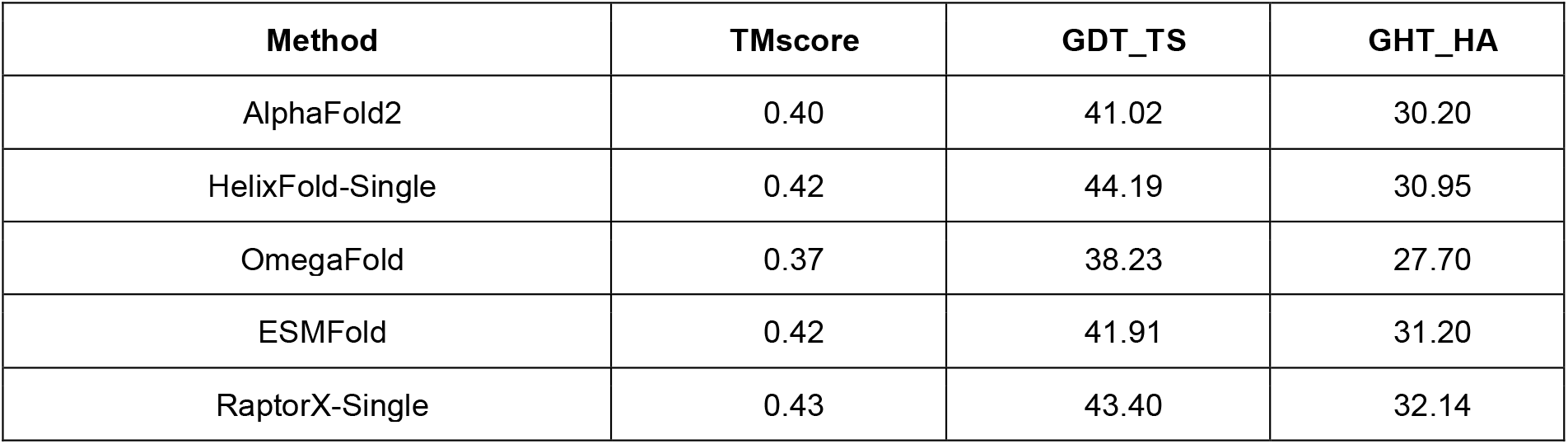
The average model quality (measured by TMscore, GDT_TS and GHT_HA) of our method, AlphaFold2, HelixFold-Single, OmegaFold and ESMFold on the Orphan dataset.

### Predicted structure changes of single mutations

Previous works showed that the MSA-based methods do not work well on predicting mutational effects [4,5,31]. To evaluate the advantage of our single sequence-based method over the MSA-based methods for mutational effect prediction, we analyze the correlation between predicted structure change and stability change of single mutations using data gathered from DMS datasets [24]. The structure change is measured by TMscore of predicted structures between the wildtype and mutated sequences. As shown in Figure 3, our single sequence methods outperform AlphaFold2 on most targets, especially on targets 1, 9 and 11. AlphaFold2 (Single) outperforms AlphaFold2 (MSA), demonstrating the advantage of single sequence methods for mutational effects prediction.

**Figure 3.**
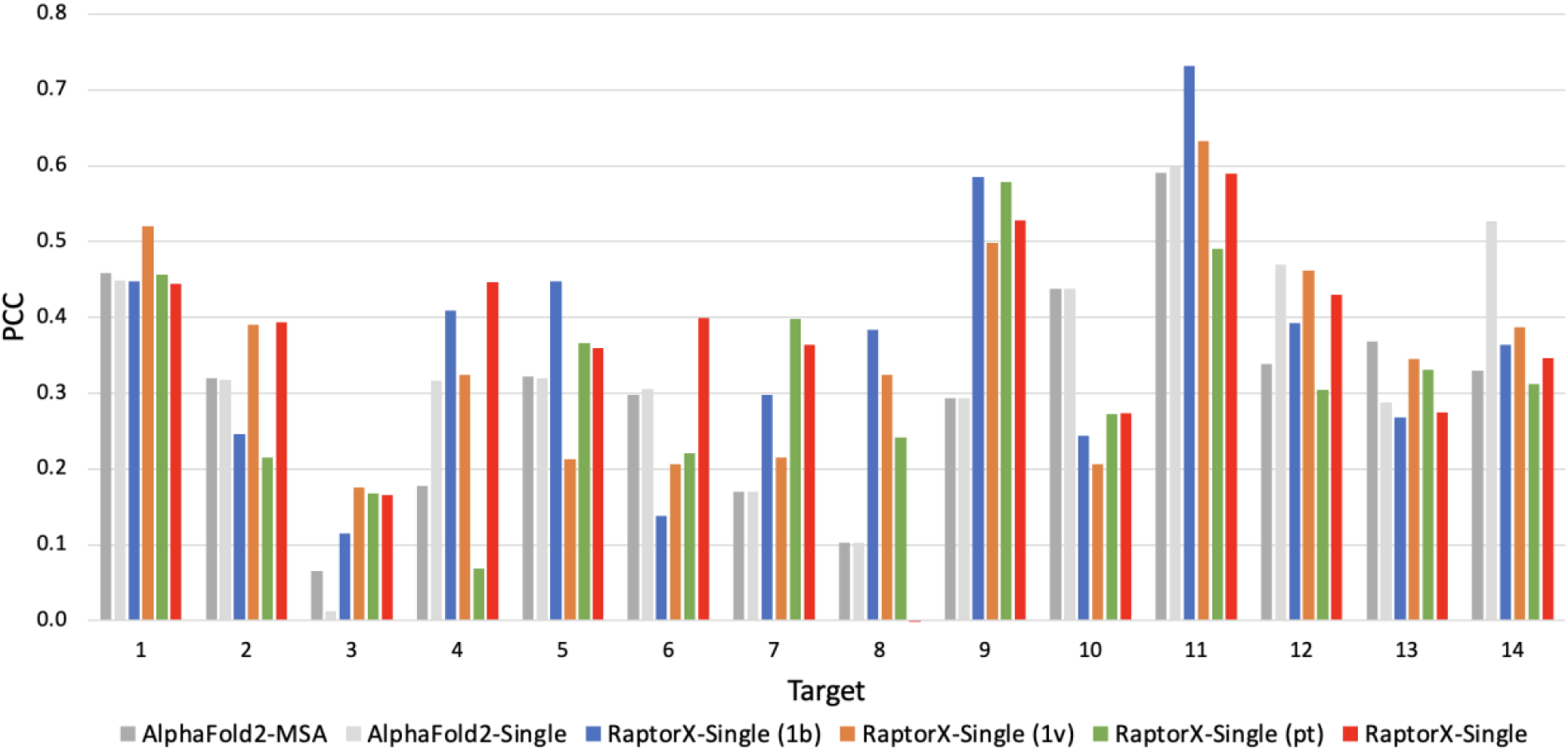
The PCC between predicted structure change and stability change of single mutations.

### Impact of MSA depth on prediction quality

It is generally believed that current single sequence-based methods are implicitly making use of sequence homologs of a protein under prediction. To study this, we compare our single sequence method RaptorX-Single with the MSA-based AlphaFold2 on targets of various MSA depths. We built a dataset consisting of 60 CASP14 targets (88 domains) and 194 CAMEO targets (released from April 23, 2022 through June 25, 2022). Most of these targets have lots of sequence homologs, to balance the MSA depth distribution, we additionally collected 99 targets released from Jan 01, 2020 through April 12, 2022 that do not have sequence homologs in Uniclust30 but may have sequence homologs in BFD, MGnify or Uniref90. As shown in Figure 4, when the MSA is shallow (<=10), RaptorX-Single outperforms the MSA-based AlphaFold2 on most targets. When the MSA is deep (>1e4), RaptorX-Single achieves comparable performance with AlphaFold2. RaptorX-Single significantly underperforms MSA-based AlphaFold2 mainly when the MSA depth is between 100 and 1000.

**Figure 4.**
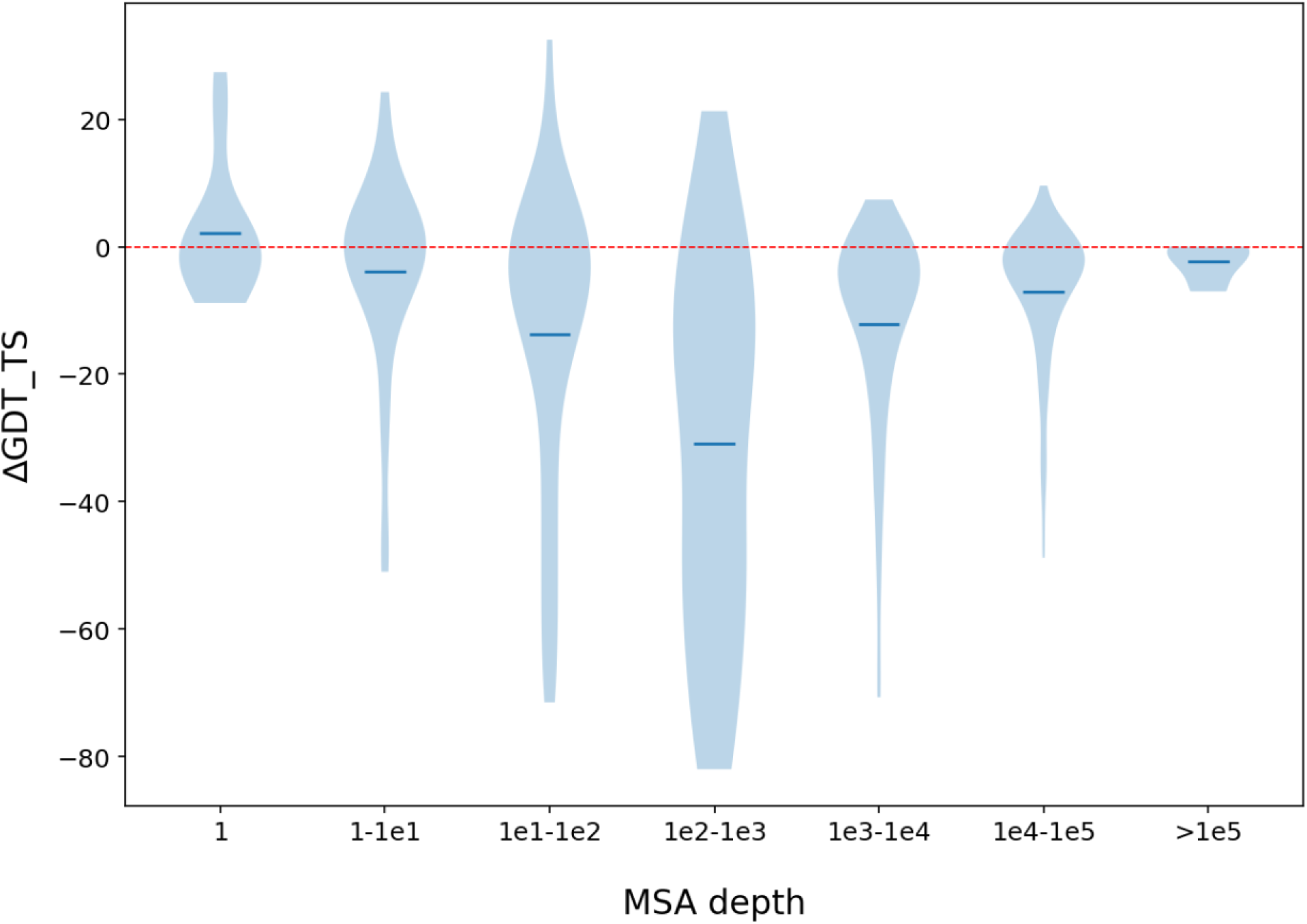
Performance comparison between our method RaptorX-Single and MSA-based AlphaFold2 with respect to MSA depth. The performance is measured by ΔGDT_TS (GDT_TS difference of the 3D models predicted by RaptorX-Single and that predicted by AlphaFold2). The MSA depth is the number of sequences in the MSA.

## Conclusion and Discussion

We have presented deep learning methods that can predict the structure of a protein without explicitly making use of its sequence homologs and MSAs. Our experimental results show that our methods outperforms MSA-based AlphaFold2 and other single sequence methods on antibody structure prediction, orphan protein structure prediction and single mutation effect prediction, demonstrating the advantage of single sequence methods on these specific structure prediction tasks. However, although single sequence methods do not explicitly make use of sequence homologs, the protein language models may implicitly encode some evolutionary and coevolution information by learning from a very large protein sequence database. Although outperforming AlphaFold2 on orphan proteins, our method and other similar ones still cannot predict the correct fold of most orphan proteins, possibly because all current single sequence-based methods are still implicitly making use of sequence homologs of a protein under prediction. Our future plan is to develop a method that can indeed predict the structure of a protein directly from its primary sequence even without implicitly using any homologous information. Such a method shall perform well on those orphan proteins.

## Author contributions

J.X. guided the research. X.J. developed the model and training code. F.W. and X.L. implemented the model training. X.J. and F.W. carried out the benchmarking. X.J. and J.X. analyzed the results and wrote the manuscript. All authors revised the manuscript.

## Acknowledgements

We are grateful to Lupeng Kong for collecting DMS data and valuable discussion.

## Availability of data

The training and fine-tuning data were derived from PDB (https://www.rcsb.org/) and SAbDab (https://opig.stats.ox.ac.uk/webapps/newsabdab/sabdab/) as mentioned in main text. The source code and benchmark target lists are available at: https://github.com/AndersJing/RaptorX-Single

## Supplementary information

**Table S1.** The average RMSD of predicted tertiary structures for the IgFold-Ab dataset.

|  | RMSD (H) |  |  |  | RMSD (L) |  |  |  |
| --- | --- | --- | --- | --- | --- | --- | --- | --- |
|  | Fr | CDR-1 | CDR-2 | CDR-3 | Fr | CDR-1 | CDR-2 | CDR-3 |
| RaptorX-Single (1b) | 0.60 | 1.31 | 1.31 | 4.61 | 0.49 | 1.42 | 0.61 | 2.32 |
| RaptorX-Single (1v) | 0.56 | 1.14 | 1.24 | 4.19 | 0.47 | 1.45 | 0.60 | 1.85 |
| RaptorX-Single (pt) | 0.55 | 1.06 | 1.17 | 4.91 | 0.47 | 1.40 | 0.64 | 2.05 |
| RaptorX-Single | 0.51 | 0.86 | 0.90 | 4.33 | 0.46 | 1.13 | 0.54 | 1.95 |
| RaptorX-Single-Ab (1b) | 0.39 | 0.67 | 0.60 | 2.79 | 0.35 | 0.76 | 0.40 | 0.92 |
| RaptorX-Single-Ab (1v) | 0.38 | 0.62 | 0.57 | 2.83 | 0.35 | 0.70 | 0.38 | 0.92 |
| RaptorX-Single-Ab (pt) | 0.38 | 0.65 | 0.60 | 2.71 | 0.34 | 0.73 | 0.38 | 0.97 |
| RaptorX-Single-Ab | 0.38 | 0.63 | 0.60 | 2.65 | 0.35 | 0.69 | 0.39 | 0.88 |

**Table S2.** The average RMSD of predicted tertiary structures for the SAbDab-Ab dataset.

|  | RMSD (H) |  |  |  | RMSD (L) |  |  |  |
| --- | --- | --- | --- | --- | --- | --- | --- | --- |
|  | Fr | CDR-1 | CDR-2 | CDR-3 | Fr | CDR-1 | CDR-2 | CDR-3 |
| RaptorX-Single (1b) | 0.72 | 1.65 | 1.41 | 5.06 | 0.65 | 1.33 | 0.82 | 2.37 |
| RaptorX-Single (1v) | 0.70 | 1.54 | 1.25 | 4.64 | 0.63 | 1.22 | 0.79 | 1.96 |
| RaptorX-Single (pt) | 0.68 | 1.40 | 1.27 | 4.89 | 0.64 | 1.23 | 0.84 | 2.25 |
| RaptorX-Single | 0.64 | 1.17 | 1.06 | 4.66 | 0.64 | 1.12 | 0.77 | 2.14 |
| RaptorX-Single-Ab (1b) | 0.57 | 1.01 | 0.83 | 3.39 | 0.53 | 0.80 | 0.66 | 1.23 |
| RaptorX-Single-Ab (1v) | 0.57 | 1.01 | 0.82 | 3.17 | 0.54 | 0.80 | 0.66 | 1.25 |
| RaptorX-Single-Ab (pt) | 0.57 | 0.99 | 0.83 | 3.32 | 0.53 | 0.81 | 0.66 | 1.25 |
| RaptorX-Single-Ab | 0.57 | 1.01 | 0.82 | 3.24 | 0.53 | 0.79 | 0.66 | 1.24 |

**Table S3.** The average RMSD of the predicted structures for the Nanobody dataset.

|  | RMSD |  |  |  |
| --- | --- | --- | --- | --- |
|  | Fr | CDR-1 | CDR-2 | CDR-3 |
| RaptorX-Single (1b) | 0.84 | 2.33 | 1.27 | 4.37 |
| RaptorX-Single (1v) | 0.81 | 2.16 | 1.18 | 4.10 |
| RaptorX-Single (pt) | 0.81 | 2.21 | 1.18 | 4.69 |
| RaptorX-Single | 0.83 | 2.19 | 1.14 | 4.06 |
| RaptorX-Single-Ab (1b) | 0.77 | 1.83 | 1.05 | 4.01 |
| RaptorX-Single-Ab (1v) | 0.73 | 1.78 | 1.10 | 3.64 |
| RaptorX-Single-Ab (pt) | 0.75 | 1.73 | 1.04 | 3.66 |
| RaptorX-Single-Ab | 0.82 | 1.78 | 1.06 | 3.50 |

**Table S4.** The average model quality (measured by TMscore, GDT_TS and GHT_HA) of our method on the Orphan dataset.

| Method | TMscore | GDT_TS | GHT_HA |
| --- | --- | --- | --- |
| RaptorX-Single (1b) | 0.41 | 42.05 | 30.34 |
| RaptorX-Single (1v) | 0.39 | 39.81 | 29.29 |
| RaptorX-Single (pt) | 0.45 | 45.25 | 33.69 |
| RaptorX-Single | 0.43 | 43.40 | 32.14 |

